# On translational control by ribosome speed in *S. cerevisiae*

**DOI:** 10.1101/513705

**Authors:** Eleanna Kazana, Tobias von der Haar

## Abstract

In addition to the widespread and well documented control of protein synthesis by translation initiation, recent evidence suggests that translation elongation can also control protein synthesis rates. One of the proposed mechanisms leading to elongation control is the interference of slow ribosome movement around the start codon with efficient translation initiation. Here we estimate the frequency with which this mode of control occurs in baker’s yeast growing in rich medium. Genome-wide data reveal that transcripts from around 20% of yeast genes show evidence of queueing ribosomes, which may be indicative of translation elongation control. Moreover, this subset of transcripts is sensitive to distinct regulatory signals compared to initiation-controlled mRNAs, and such distinct regulation occurs for example during the response to osmotic stress.

**Note:** the previous version 2 of this preprint contained a Decision-Tree based analysis where we attempted to relate mRNA features to the presence or absence of queueing ribosome peaks. Since releasing that version, we performed additional controls for this analysis which strongly sugest that its results are random and should be ignored. Specifically, both the overall predictability and the importance of individual features is very similar for a real dataset where genes are labelled as containing a second SSU peak or not, and for a simulated control dataset containing an equal proportion of randomly labelled genes. We have removed this figure from the current version. The analysis together with the additional control remains accessible on the accompanying Github repository (github.com/tobiasvonderhaar/ribosomespeedcontrol) in the file “Figure 3 Obsolete (Classification).ipynb”.

## Introduction

Translation exerts control over gene expression, both in terms of setting the basal protein production rate for an mRNA and in terms of adapting production rates to challenging environments or to developmental needs (1). The three stages of translation (initiation, elongation and termination) make distinct contributions to translational control. Translation initiation factors, which mediate the formation of productive ribosome-mRNA contacts, were long thought to be the more or less exclusive targets for translational control. However, recent studies have revealed that translation elongation, ie the movement of ribosomes along the ORF and the concurrent tRNA-dependent decoding of the codon sequence, can also be targeted. For example, elongation can be rate limiting in cancers (2), dynamic tRNA modifications (3) and regulation of translation elongation factor 2 by phosphorylation (4) can exert translational control over individual transcripts by modifying elongation rates, and the regulation of translation elongation during cooling (5) mediates effects of sub-physiological temperatures (6).

While it is clear that translation elongation can control gene expression levels, the molecular mechanisms by which this happens are not comprehensively understood. Two specific mechanisms that have been identified which can connect translation elongation rates to expression levels is the Dhh1-dependent destabilisation of mRNAs containing slowly decoded codons (7–9), and interference of slow moving ribosomes with efficient translation initiation (10). The relative importance of these two mechanisms for different genes has not been studied on a genome-wide basis, although our previous study on four recombinant proteins suggests that both predominantly translational regulation-mediated and predominantly mRNA stability-mediated regulation can coexist in different transcripts in the same cell (cf. figure 2 in ref. (10).

**Figure 1.**
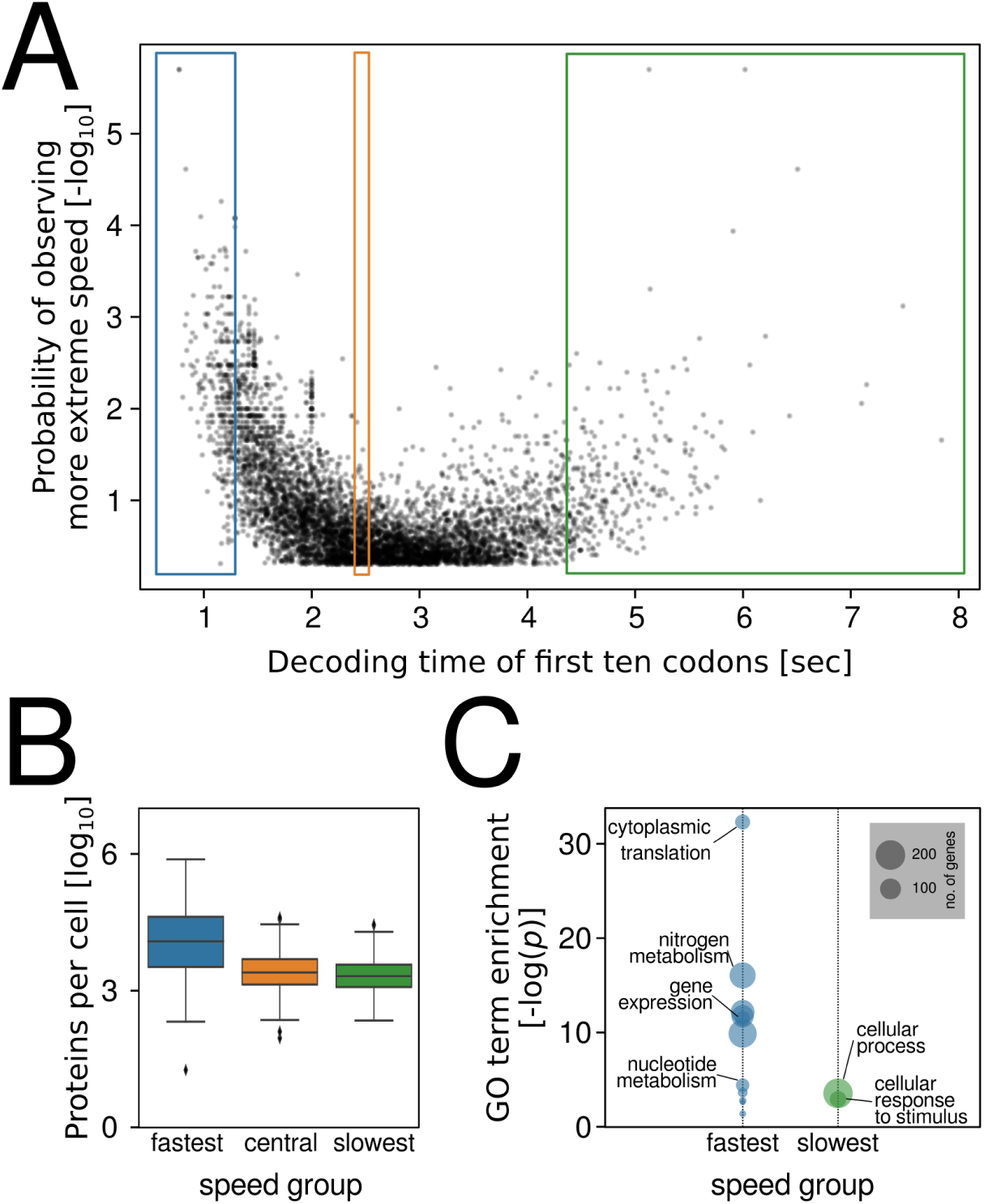
AUG-proximal speed throughout the yeast genome. **A**, the decoding speed of the first ten codons of each yeast ORF was estimated based on our published computational models of codon decoding (10), and compared to the distribution of possible speeds for the same protein sequence. The plot relates the observed decoding time with the log probability of observing a more extreme decoding time. The coloured boxes enclose data points analysed in panel B. **B**, protein abundance of genes highlighted in panel A. **C**, GO annotation enrichment for genes highlighted in panel A.

**Figure 2.**
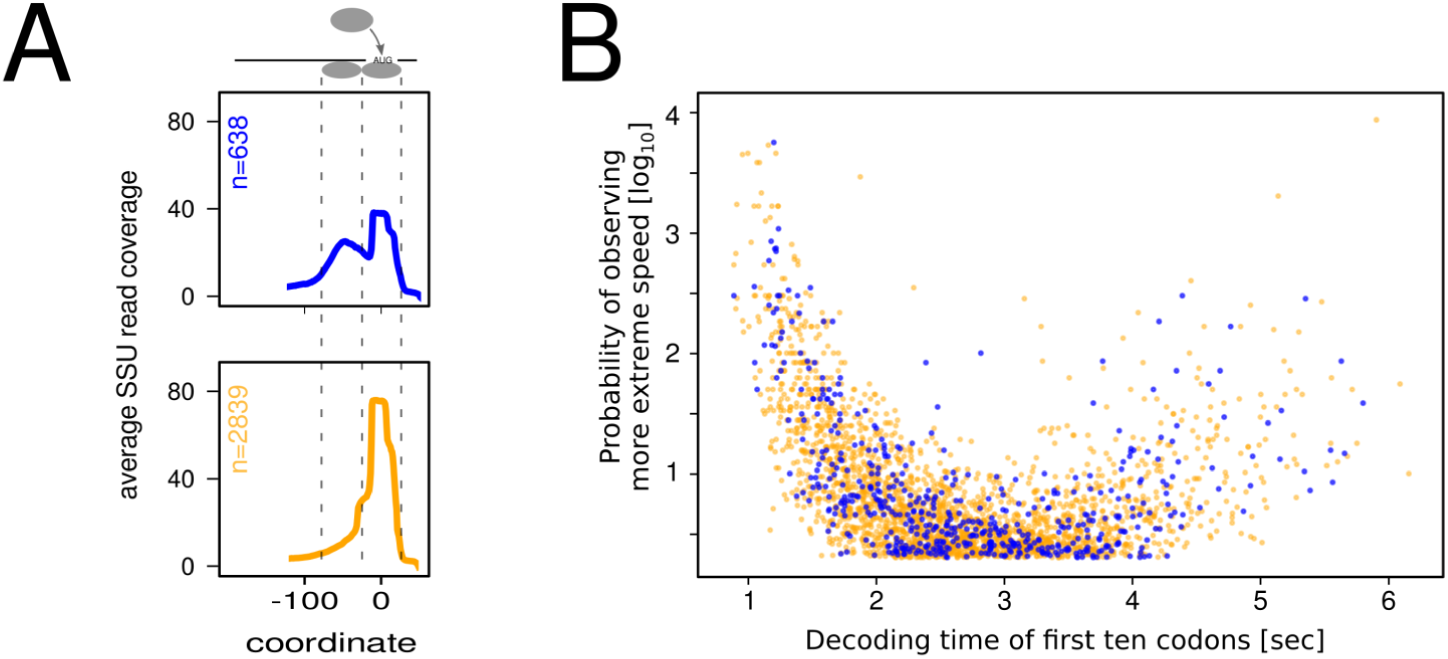
Scanning queues throughout the yeast genome. **A**, metagene plots of small ribosomal subunit footprinting data from Archer *et al*. (19), for genes with 5’-UTR lengths longer than 45 nucleotides. Separate metagene plots are shown for two classes of foot-prints, classified by the presence or absence of a second peak upstream of the main AUG peak corresponding to initiating subunits. Metagene plots are aligned to the start codon at position zero. **B**, mapping of genes displaying signals for waiting small ribosomal subunits throughout the yeast genome. Colours correspond to panel A. The volcano pot is identical to figure 1A, but only shows genes with 5’-UTR lengths above 45 nucleotides.

The identification of individual translation elongation-controlled mRNAs has so far been mostly anecdotal, (6, 11, 12). Here, we sought to systematically identify translation elongation-controlled genes in baker’s yeast, focussing specifically on such genes where elongation control is mediated at the level of translational efficiency, rather than Dhh1-dependent mRNA stability effects. We find that around 20% of yeast genes display signals in ribosome footprinting experiments that may be consistent with translation elongation control, and for a small selection of these genes we provide experimental evidence that enhancing elongation speed does indeed increase their translation rates. We show that this subset of the transcriptome can be controlled by distinct regulatory signals, and that this may lead to distinct response dynamics during the onset of stresses.

## Materials and Methods

### Analyses and data availability

Unless otherwise stated, data analyses were conducted using Python 3.0. Feature selection was performed using Scikit-learn (13), and statistical analyses were performed using scipy.stats (14). All analysis scripts, Literature datasets and experimental raw data are available on GitHub.

### Data sources

Protein expression levels were retrieved from a recent meta-analysis of protein abundance data by Ho *et al*. (15). 5’-UTR length data were retrieved from the supplemental material from a number of studies (16–18). The longest reported 5’-UTR value reported in any of these three studies was used to apply the 45-nucleotide cut-off for analysis of the SSU footprinting data. The SSU footprinting data published by Archer *et al*. (19) were retrieved using the table browser function of GWIPS-Wiz (20). uORF data were retrieved from Ingolia *et al*. (21). Data on secondary structure content were from Kertesz et al. (22) and translation initiation rates from Ciandrini *et al*. (23).

### Yeast strains and plasmids

The standard yeast strain used in this work was BY4741 (24). All gene deletions were in this background and were from the systematic deletion collection (25) except for the *tef1::HIS3* deletion strain which was a kind gift from Paula Ludovico (University of Minho, Portugal).

Plasmids are listed in table 1. Plasmid DNA, detailed plasmid maps and sequences are available through the Addgene repository.

**Table 1.** Plasmids used in this study.

|  |  |  | Reference | Addgene No. |
| --- | --- | --- | --- | --- |
| pTH825 | Reporter I | Standard codon usage firefly luciferase preceded by a uORF-containing 5'-UTR, codon disoptimised *Renilla* luciferase preceded by a short 5'-UTR | This study | 115370 |
| pTH862 | Reporter II | Codon optimised *Renilla* luciferase preceded by a uORF-containing 5'-UTR, codon disoptimised firefly luciferase preceded by a short 5'-UTR | This study | 115371 |
| pTH727 | Reporter C | Standard codon usage *Renilla* and Firefly luciferase genes preceded by short 5'-UTRs | (10) | 38211 |

<sup>1</sup> github.com/tobiasvonderhaar/ribosomespeedcontrol
<sup>2</sup> www.addgene.com

pTH825 (Reporter I) was generated by replacing the *Renilla* luciferase (RLuc) gene in pTH743 (10) with a codon-disoptimised RLuc gene described in the same publication, using *XmaI* and *EcoRI* restriction sites introduced adjacent to the start and stop codon by PCR.

pTH862 (Reporter II) was generated by cloning a *Renilla* luciferase gene which had been codon optimised as described (10) into pTH644 (26) using *XmaI* and *EcoRI* sites. A Gcn4-derived, uORF containing 5’-UTR sequence was amplified from pTH743 and introduced into the *XmaI* site preceding the codon-optimised RLuc gene. Finally, the firefly luciferase gene from pTH726 (10) was introduced into the *BamHI* and *HindIII* sites of the new vector.

### Gene replacement strains

A strain containing the codon optimised *HIS3* gene has been described (10).

To compare expression levels of the wild-type and optimised *SUP35* gene, the optimised ORF sequence was combined with the natural *SUP35* promoter and terminator sequences in plasmid pUKC1620 (27) using a Gibson assembly strategy (28). The resulting plasmid and a wild-type plasmid for comparison were shuffled into yeast strain LJ14, which contains a chromosomal deletion of the *SUP35* gene (27).

For the five remaining genes analysed in this study, the general strategy for constructing the gene replacement strains included the following steps: 1) design and synthesis of codon optimised sequences including upstream and downstream flanking sequences to facilitate homologous recombination with the corresponding genomic locus; 2) generation of CRISPR guide RNA vectors targeting the gene; 3) co-transformation of a wild type yeast strain with a guide RNA vector and the matching linearised, optimised gene; 4) confirmation of integration of the optimised gene by diagnostic PCR; and 5) assessing resulting changes to protein and RNA levels using western blotting and qPCR. Detailed procedures for these steps are given in supplemental file 2.

### Dual luciferase assays

These were conducted in 96-well format as described (29). Prior to analyses, plates containing the source cultures were visually inspected for contaminated wells, and corresponding data points were disregarded for data analyses.

### Western Blots

Protein extracts were prepared and western blots performed as described (30). Rabbit antibodies were sourced from the following publications or companies: anti-HA (Sigma Aldrich, UK, H6908), anti-Cdc10 (Abmart, NJ, USA, X2-P25342), anti-Ras2 (santa Cruz Biotechnology, TX, USA, Sc-6759), anti-Sup35 (27), anti-Ade2 (31), anti-Grx5 (32), anti-NBP35 (33).

## Results

### 5’-proximal ribosome speed throughout the yeast transcriptome

Because start codons can only be occupied by a single ribosome at a time, mRNA-specific protein synthesis rates are controlled by translation elongation whenever the rate of translation initiation attempts exceeds the rate with which ribosomes elongate away from the start codon (10). The rate of ribosome movement is sequence dependent and controlled by codon usage patterns (34), mRNA secondary structure (35), the charge of the nascent chain within the ribosomal exit tunnel (36), and the readiness of particular amino acids to undergo peptidyl transfer (37). Although ribosome speed can thus be affected by many different mechanisms, models which exclusively consider tRNA:codon interactions as determinants of ribosome speed can predict protein yields with high degrees of accuracy (10, 26). This, together with the recent finding that ribosome collision sites are strongly enriched near the start codon (38), predicts that codon-dependent ribosome speed near the start codon is an important determinant of translational control. We therefore initiated our analyses of elongation-controlled mRNAs in the yeast transcriptome by analysing codon-dependent ribosome speed immediately following the start codon, as a non-exhaustive but useful potential indicator of mRNAs which might be subject to control by translation elongation.

We used our published, model-based decoding time estimates for each codon (10) to calculate the speed of decoding of the first ten codons following the start codon (approximately the span of one ribosome) throughout the entire yeast transcriptome. In addition to the decoding speed of the observed sequences, we also computed decoding speeds of random sequences encoding the same peptides as the actual genes, and determined the proportion of sequences that were more extreme in their speed than the observed sequence. The negative logarithm of this proportion (*P*) gives an estimate of the probability of observing the actual decoding speed of a gene if codon usage for this gene was entirely random.

Plotting *P* against absolute speed for all yeast genes results in a volcano plot (figure 1A) which displays a clear skew towards fast sequences, reflecting the well-documented general codon bias in the yeast genome (39). To explore the observed distribution further, we selected the fastest, slowest, and central 5% of genes and analysed these groups with respect to particular features. The fastest group was associated with significantly higher expression levels than the other two groups, and the very highest expressed proteins were found exclusively within this group (figure 1B). The fastest group also showed strong association with specific GO terms including cytoplasmic translation, consistent with the observation that ribosomal proteins show particularly high codon usage bias (40). In contrast, the slowest group of genes was not associated with lower expression levels compared to the central group and was much less strongly associated with specific GO terms. This indicates that fast sequences are favoured in the yeast genome through evolutionary selection on genes whose functions require particularly high expression. In contrast, we detected no significant selection for slowly decoded sequences.

This observation is relevant to the question how and why elongation-controlled mRNAs arose in the yeast genome. A feature that places an mRNA under control of translation elongation is a combination of (relatively) high initiation and (relatively) slow elongation rates. In a population of mRNAs where initiation and elongation rates evolve randomly and independently, combinations of fast initiation and slow elongation could arise simply by chance. In this case, elongation-controlled mRNAs would be enriched among those that contain particularly slow codons, where the probability of a high initiation/elongation rate ratio is highest. Our observation that such mRNAs are not widely selected for implies that either elongation-controlled mRNAs themselves are also not widely selected for, or that selection for elongation-controlled mRNAs occurs by combined selection on initiation- and elongation-rate determining features.

### Characterisation of the elongation-controlled transcriptome

In recent work Archer *et al*. (19) used translation complex profiling, a variant of the ribosome profiling approch (41), to study footprints on mRNAs derived solely from small ribosomal subunits (SSUs). Since our previous findings indicate that mRNAs become elongation controlled when initiating ribosomes are prevented from accessing the start codon because the previous ribosome has not yet liberated this site (10), we expected that scanning 40S subunits may form queues 5’ of the start codon on such mRNAs. Queueing 40S subunits should be detectable in the Archer *et al*. dataset. If we assume that initiating ribosomes physically cover around 30 nucleotides centred around the start codon itself, the centre of any small ribosomal subunit queueing immediately upstream should be somewhere in the region spanning nucleotides -60 to -15 upstream of the AUG (note that due to the trailing mass of translation initiation factors, scanning 40S subunits produce larger footprints with less clearly defined boundaries compared to elongating ribosomes (19)).

We retrieved the footprinting data generated by Archer *et al*. from the GWIPS-viz database (20), processed the data for each gene using a peak calling algorithm, and classified the genes into those containing an identifiable second peak of SSU footprints within the -60 to -15 region adjacent to the main SSU footprint peak over the ORF start codon, and those without such a peak (figure 2A). Because upstream footprints can only occur on sufficiently long 5’-UTRs, we restricted this analysis to those mRNAs having 5’-UTRs longer than 45 nucleotides. All of the 3477 genes with a 5’-UTR length above this cut-off showed a detectable small subunit peak centering round the start codon. In addition, 638 (18.3%) of these genes also showed a detectable second upstream peak, with a mean peak location around 50 nucleotides from the start codon (figure 2A).

We propose that the mRNAs displaying a second 40S peak correspond to the elongation-controlled transcriptome (further experimental evidence for this is given below). However, our analyses could be confounded by other mechanisms attracting small ribosomal subunits to sites upstream of the main start codon, notably by translation initiation events on upstream open reading frames (uORFs). We therefore analysed the relationship between uORFs and apparent queuing SSU peaks in more detail. The proportion of genes with uORFs is similar in the gene subsets with and without a second peak (p=0.67 by Fisher’s Exact Test), and second SSU peaks are therefore not generally associated with uORFs. Of those genes displaying second SSU peaks and also containing uORFs, the majority of uORF locations is outside of the -15 to -60 nt analysis window, and the observed second footprint peaks can thus not be results of uORF initiation (supplemental figure 1A). Genes which do have uORFs within the analysis window comprise less than 5% of the “second SSU” set. Spot checks with representative genes (supplemental figures 1 B and C) indicate that for some but not all of these the second SSU footprints may indeed arise from uORF initiation events rather than from ribosome queuing, in particular those where the main ORF AUG footprints are substantially lower than the queuing footprints. However, overall this suggests the proportion of genes mis-annotated as subunit queuing genes in our analyses due to interference from uORFs is << 5%.

When mapped against the 5’-decoding speed of the yeast transcriptome, the corresponding mRNAs appear randomly distributed without any strong association with specific speed properties (figure 2B). Based on the considerations outlined in the previous section, this suggests that such mRNAs may have arisen as the result of selection on particular initiation/elongation rate ratios, rather than selection solely for slow movement. To explicitly test the assumption that mRNAs displaying a second peak in the Archer *et al*. dataset correspond to elongation-controlled mRNAs, we manipulated the ribosomal decoding speed of a number of yeast genes at their normal chromosomal loci *in vivo*. This involved designing and synthesizing speed-optimised gene sequences, replacing the original genes in the yeast genome using a CRISPR-based approach, and finally assessing the effect of speed optimisation on mRNA and protein expression levels for this gene. For any gene where ribosome speed restricts achievable translation initiation rates, we expect increases in decoding speed to increase the protein/mRNA ratio, but this should not be the case for initiation-controlled genes.

This assay, which we had originally used to demonstrate elongation control of the yeast *HIS3* gene (10), is laborious and cannot be applied to large numbers of genes, but we reasoned that it would be a good way of verifying predictions from the Archer *et al*. dataset using a smaller number of genes. In addition to the *HIS3* gene, we applied the assay here to six additional genes, for which we could source high quality antibodies and which span a range of decoding speeds (figure 3A). Importantly, these genes were selected before the results from the SSU foot-printing analyses had been completed, and the experimenters were blind to the results from these analyses throughout the experimental procedure.

The results of these assays are displayed in figure 3B-H. The *HIS3* gene (figure 3B) has a very short 5’-UTR of 10 nt (17), which is too short to accommodate a queueing 40S subunit. Although the protein/mRNA ratio for the His3 protein is increased by speed optimising the gene, confirming that the mRNA is elongation-controlled, this gene therefore does not show a second SSU peak in the Archer *et al*. dataset. The remaining six genes all have 5’-UTR lengths that should permit queueing ribosomes to form footprints, but only one of these (*RAS2*) actually showed clear evidence of a second footprint. *RAS2* is also the only one of the six additionally tested genes for which the replacement assay led to a clear increase in the protein/mRNA ratio, confirming the link between elongation control of protein abundance and the presence of an upstream SSU footprint. *CDC10* showed a weak increase in the protein/ mRNA ratio as well as displaying a weak queueing ribosome peak, however these signals were obscured by the very low expression levels for this gene. The other four genes neither showed increased mRNA/protein ratios, nor evidence of upstream SSU footprints. Thus, for the sample of genes tested with this assay, the codon replacement assay and the SSU footprinting dataset arrive at a unanimous classification of genes as either elongation or initiation controlled. The observation that 18% of yeast genes with sufficiently long 5’-UTRs show footprints for queueing small ribosomal subunits thus suggests that this number is a good first estimate for the proportion of elongation-controlled genes in the yeast genome generally.

It should be noted that this proportion is specific to the physiological conditions of our assays, ie logarithmic growth in rich medium. Any change in growth conditions that entails, for example, a specific reduction in translation elongation rates would lead to the transfer of additional mRNAs into the elongation-controlled pool and vice versa. Under conditions of strong regulation (eg when translation initiation is essentially halted by phosphorylation of eIF2 during amino acid starvation, or when translation elongation is stalled globally by activation of eEF2 kinases) all mRNAs should become controlled by either translation initiation or elongation.

### Separable regulation of initiation- and elongation-controlled transcripts

While *substantial* regulation of translation factor activity is predicted to eventually affect all transcripts, we reasoned that *limited* regulation of initiation factor activity should preferentially affect initiation-controlled mRNAs, whereas limited regulation of elongation factors should preferentially affect elongation-controlled mRNAs. The coexistence of initiation- and elongation-controlled sets of mRNAs in a cell could thus split the transcriptome into two separately addressable regulons.

To test the concept that such separate regulation can indeed occur, we designed a series of reporter constructs expressing two luciferases (figure 4A). mRNAs encoding the two luciferases were placed either under initiation control (by combining an inefficient, uORF-containing 5’-UTR with an efficient, codon-optimised ORF), or under elongation control (by combining an efficient 5’-UTR with an inefficient, slow-codon containing ORF). In construct I (figure 4A), we paired an initiation-controlled firefly luciferase with an elongation-controlled *Renilla-*luciferase gene, whereas the control regimes were inverted in construct II. In a control construct, C, both luciferases were placed under elongation control. By normalising the expression ratios observed with constructs I and II to construct C, and by ensuring that constructs I and II changed expression in opposite ways, we could reliably separate changes in expression resulting from the regulation of translation initiation or elongation from expression changes resulting from other regulatory events including transcription and protein turnover.

**Figure 4.**
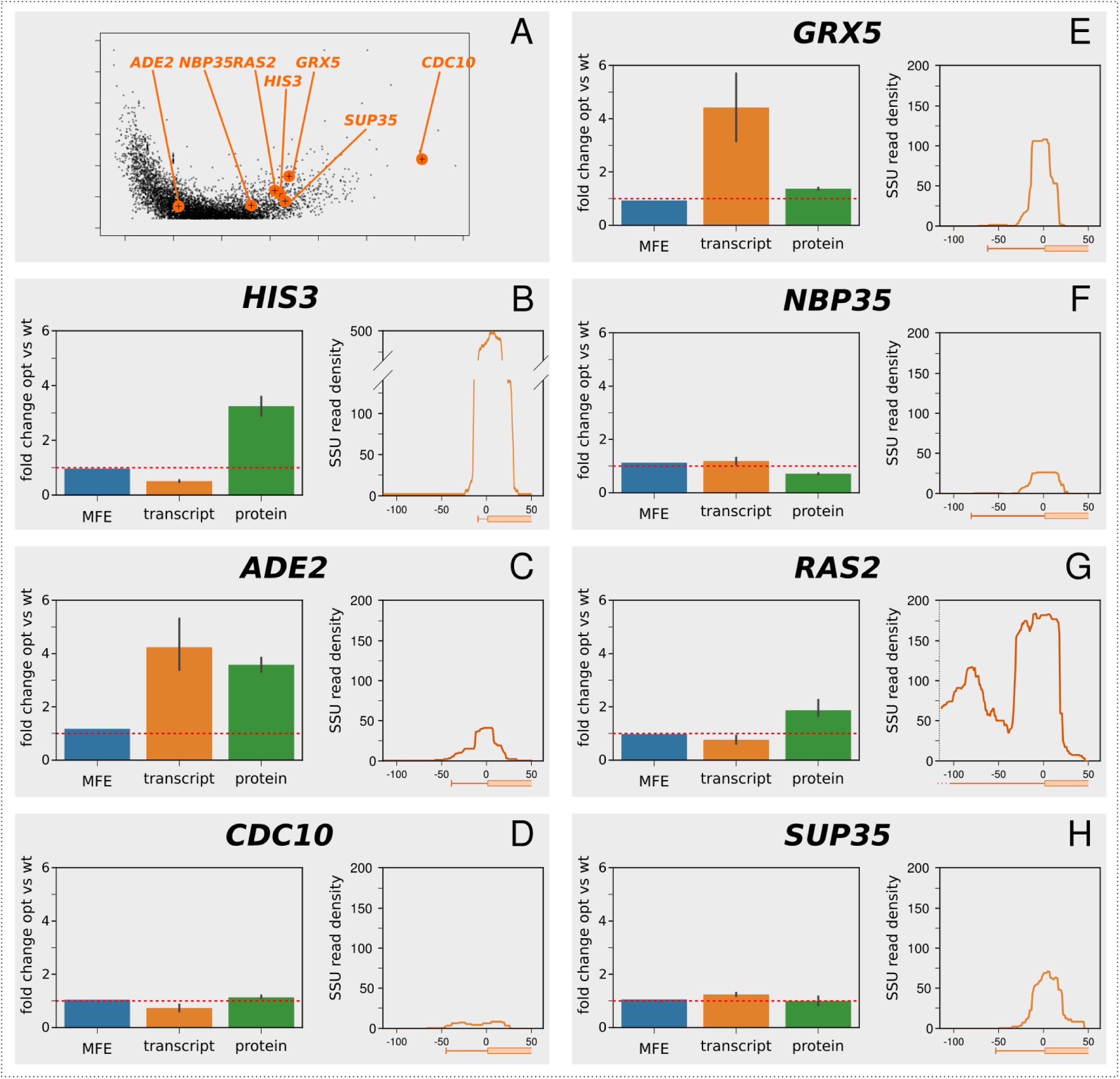
Control analysis of selected genes. **A**, the location of the selected genes is indicated in the volcano plot from figure 1A. **B-H**, analyses of individual genes. For each gene, the predicted change in folding energy (as the mean free energy of the ensemble, MFE), the measured change in mRNA abundance, and the measured change in protein abundance are shown for the codon optimised vs the original gene. The red lines indicate a ratio of 1 ie where codon optimisation does not change the quantified parameter.

We measured changes in the expression ratio of the two luciferases in yeast strains containing a deletion of one of two identical isogenes for translation initiation factor 4A (*tif1Δ*, mimicking limited regulation of translation initiation), or a deletion of one of two identical isogenes for translation elongation factor 1A (*tef1Δ*, mimicking limited regulation of translation elongation). We observed that in the *tif1Δ* strain, where initiation activity was reduced, the expression ratio changed in favour of the elongation-controlled luciferase in both constructs I and II (figure 4B). In contrast, in the *tef1Δ* strain, the expression ratio changed in favour of the initiation-controlled luciferase in each construct. We further tested the behaviour of our constructs upon addition of the translational inhibitor, cycloheximide. This inhibitor is generally used at high concentrations as a translation elongation inhibitor that interferes with the translocation step (43), although at low concentrations *in vivo* it is thought to act as a translation initiation inhibitor before inhibiting elongation at higher concentrations (44). In our reporter constructs, the ratios of the two luciferases changed with cycloheximide addition in a concentration dependent manner consistent with this notion of dual inhibition. At low concentrations, the elongation-controlled luciferases were initially favoured but this trend became reversed at higher concentrations (figure 4C). Thus, both genetic and chemical manipulation of translation initiation and elongation rates indicated that transcripts under distinct control regimes can indeed be separately addressed by translational control mechanisms.

### Initiation- and elongation-control during stresses

Having shown that initiation- and elongation-controlled yeast transcripts could in principle be addressed via distinct regulatory mechanisms, we wished to explore in how far this was used during natural gene expression regulation. For this purpose, we measured the luciferase ratios of our reporter assays under a number of different growth and stress conditions. We observed conspicuous divergent shifts of the luciferase ratios with changes in temperature (figure 5A), which were consistent with relatively slower elongation rates at lower temperatures, and increased elongation rates at higher temperatures. Although this is to our knowledge the first report of regulation of translation elongation in sub-physiological temperatures in yeast, these findings mirror the known regulation of translation elongation during cooling in mammalian cells (5) and tissues (6).

**Figure 5.**
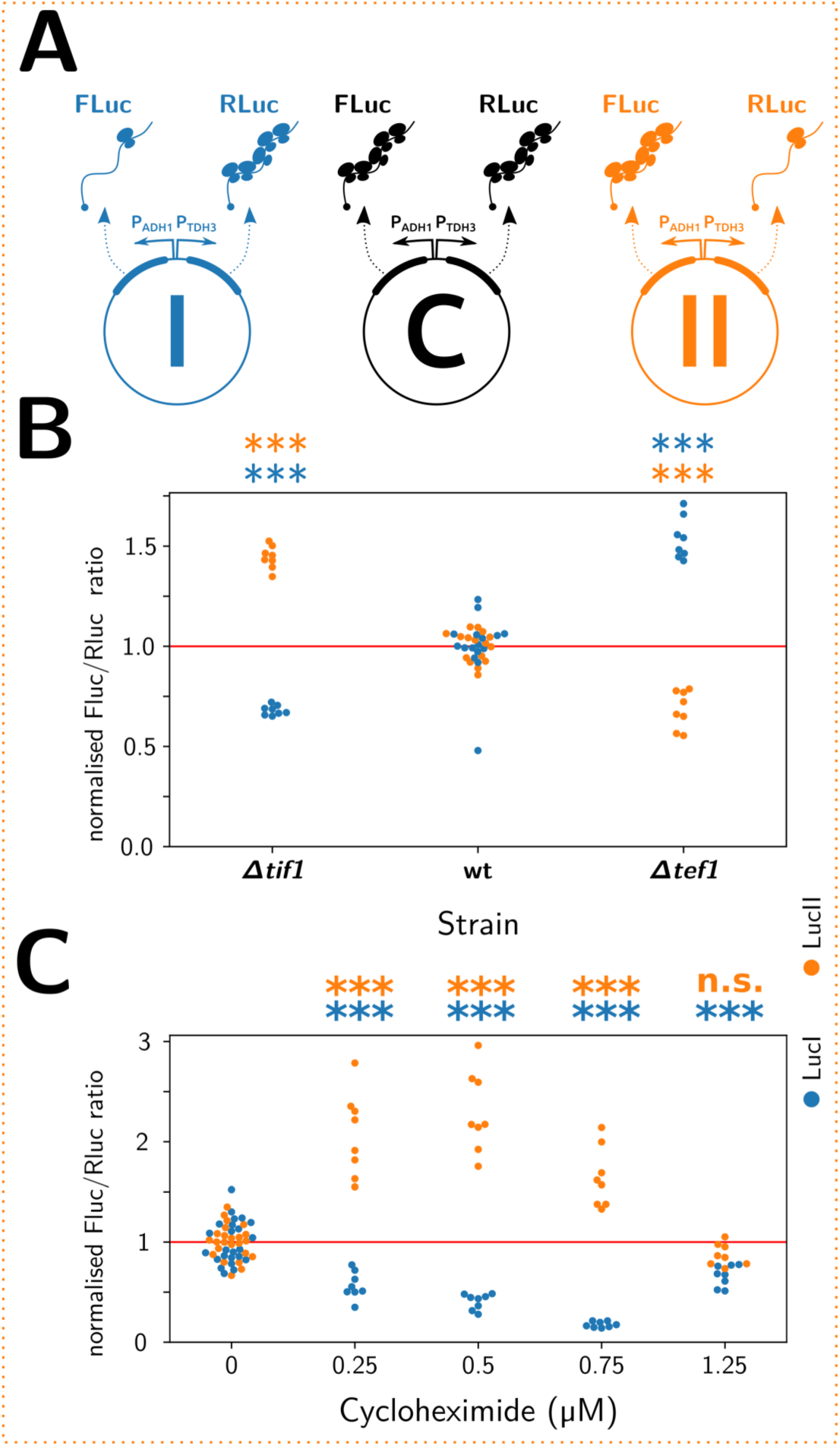
Elongation- and initiation-controlled mRNAs have distinct regulatory properties. **A**, a series of reporter plasmids containing separately assayable luciferase genes either under initiation-control (FLuc in repoter I and RLuc in reporter II) or under elongation-control (Rluc in reporter I and FLuc in reporter II). Reporter C is designed to control for changes in transcriptional and post-translational regulation, and all data are normalised against the FLuc/RLuc ratio of this reporter. **B**, strains which contain moderate translation initiation defects (*Δtif1*) or moderate translation elongation defects (*Δtef1*) shift the expression ratio between the luciferases, favouring expression of the mRNA controlled by the non-deficient pathway. **C**, upon application of various concentrations of cycloheximide, which acts as a translation initiation inhibitor at low concentrations but as an elongation inhibitor at high concentrations, the expression ratios initially favour the elongation-controlled reporter before returning to a neutral ratio. Significance of the difference to the control condition (samples “wt” in panel B and “0” in panel C) was determined by ANOVA and Tukey’s test and is indicated (***, p<0.001; n.s., p>0.05).

We also tested a number of stress conditions including oxidative, osmotic and cell wall stresses. Most of these did not affect the expression balance between the initiation- and elongation-controlled reporter genes, with the conspicuous exception of the two osmotic stress conditions we tested (0.5 M NaCl and 1 M sorbitol, figure 5B). For both of these conditions, the observed expression pattern was consistent with a clear initiation block. The response to osmotic stress has been studied in detail by a number of authors and this response is known to involve regulation at both the levels of translation elongation (45) and initiation (46). The elongation response, which involves activation of the Hog1 kinase, is known to be transient persisting for less than 10 minutes under a low intensity stress (0.2 M NaCl, (47)) and for around 30 minutes for a stronger stress (0.8 M NaCl, (48)). The dynamics of the regulation of initiation has not been studied in detail but it is known that this component is important for re-establishing translation following the initial sharp downregulation upon onset of the stress. Since the measurements in figure 5B are performed under steady-state stress conditions (ie after the initial, transient regulatory events have passed), these findings indicate that an initial translational arrest due predominantly to the inhibition of translation elongation is followed by a steady-state response in which translation elongation recovers, but translation initiation is partially inhibited compared to pre-stress conditions.

We sought to more closely examine the responses of initiation- and elongation-controlled mRNAs during the initial, elongation-regulating response to osmotic stress conditions. We attempted to measure the luciferase ratios from our reporter constructs, but found that due to different protein half-lives of the firefly and *Renilla* luciferases this assay is not suitable for application outside of steady-state conditions (data not shown). However, Lee *et al*. generated detailed, genome-wide mRNA and protein abundance data at timepoints immediately following the onset of a 0.7 M NaCl stress (49), and we used these data to study the evolution of protein/ mRNA ratios over time following the onset of the stress. For this analysis, we initially split the yeast transcriptome into elongation- and initiation-controlled sets as shown in figure 2A (again only using the subset of mRNAs with 5’-UTR lengths >45 nt, for which footprints of queueing SSUs are observable). We then clustered each of the two sets according to the time-evolution of the protein/ mRNA ratios, in order to determine subsets of mRNAs that showed similar behaviour (figure 6). For a cluster number of four, this analysis revealed qualitatively very similar clusters for both initiation- and elongation-controlled mRNAs. However, although the behaviour was qualitatively similar, a quantitative comparison showed that, on average, within each cluster the elongation controlled mRNAs were more repressed at the initial time point (t=10 minutes) than the initiation controlled mRNAs. At this time point, the Hog1 response is known to be maximal (48). For all four clusters, this difference at the initial time point was statistically significant (figure 6). Overall, these analyses showed that under a stress condition where the regulation of translation elongation makes a strong contribution to the cellular response, the initiation- and elongation-controlled subsets of the transcriptome respond with distinct dynamics.

**Figure 6.**
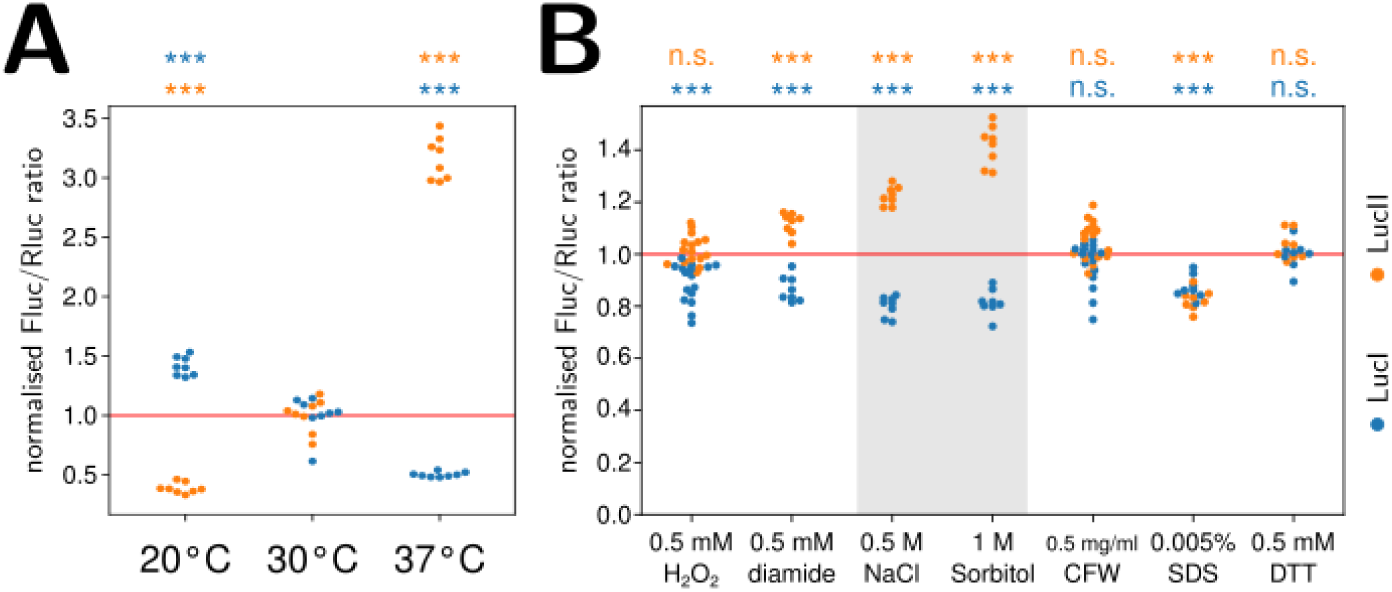
Distinct elongation- and initiation-control under stress conditions in yeast. **A**, changes in temperature shift the expression ratio of the reporter RNAs consistent with an elongation block at lower temperatures. **B**, some stress conditions shift expression ratios of the reporter mRNAs consistent with distinct regulation of translation initiation and elongation pathways. This appears particularly strong for osmotic stresses (0.5 M NaCl and 1 M sorbitol, shaded). Significance of the difference to the control condition (sample “30°C” in panel A and non-supplemented medium in panel B) was determined by ANOVA and Tukey’s test and is indicated (***, p<0.001; n.s., p>0.05).

**Figure 7.**
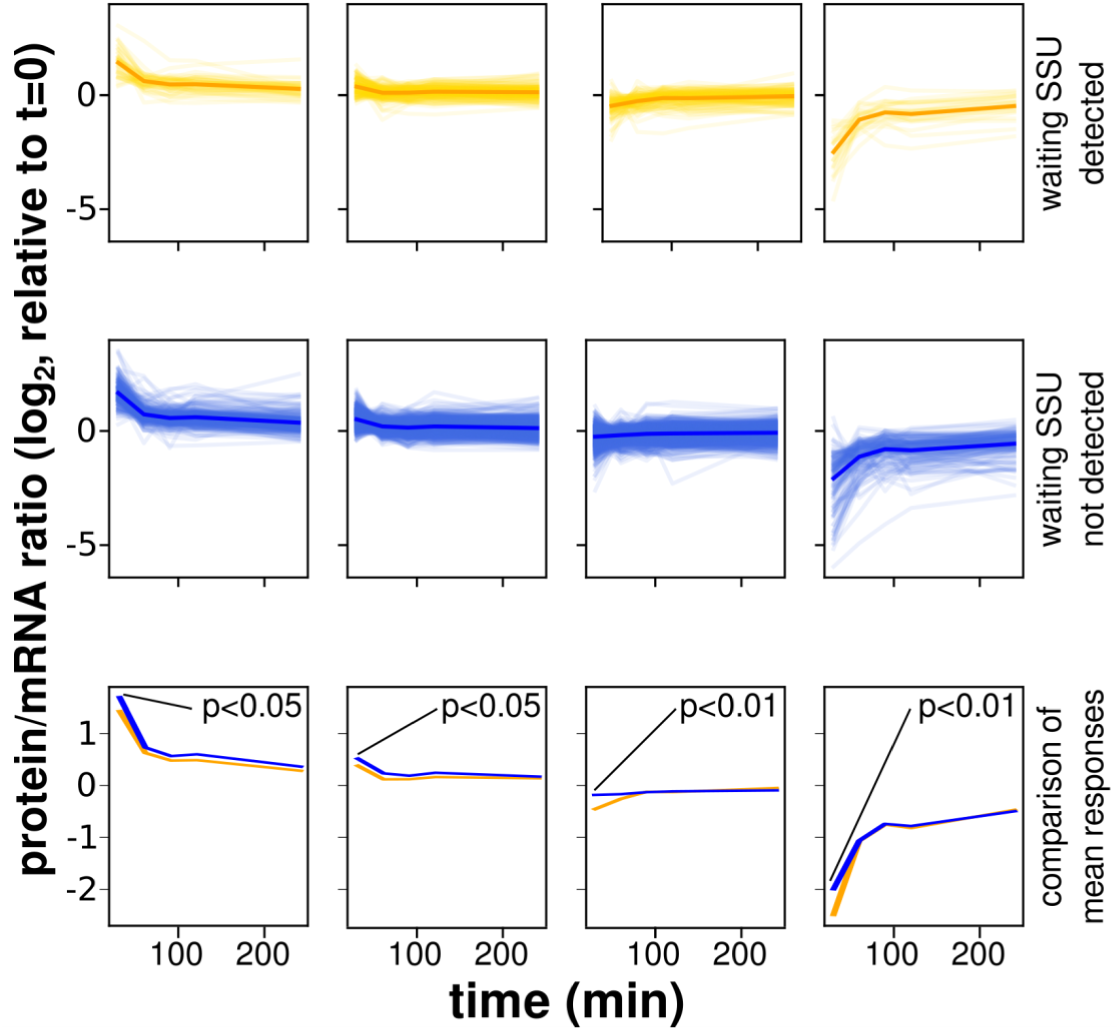
Distinct response dynamics of initiation- and elongation-controlled mRNAs under conditions known to entail regulation of translation elongation. These analyses are based on a dataset published by Lee *et al*. (49), which reported the evolution of protein and mRNA abundances following the acute onset of an osmotic stress. All genes with 5’-UTR lengths >45 nt were divided into those containing evidence of queueing small ribosomal subunits (yellow) or no such evidence (blue). Genes were then clustered into classes where the protein/mRNA ratio followed similar time evolution. In the top two rows, faint traces correspond to individual genes whereas solid traces are metagene plots for all genes in that same plot. In the bottom row, the metagene traces of the plots above are replotted to facilitate comparison. In all classes, mRNAs with queueing subunits show reduced protein:mRNA ratios at the initial time-point where activation of the Hog1 kinase, which suppresses elongation factor 2 activity, is known to be strongest. Significance of the difference for the timepoint at t=10 minutes is indicated.

## Discussion

The translational control field has historically focussed on the role of translation initiation factors in regulating protein synthesis efficiency (1). Following several anecdotal descriptions of individual mRNAs from various organisms for which elongation was instead found to be rate limiting (6, 10, 11, 50), we sought to determine how widespread this mode of control is in the model eukaryote, baker’s yeast. We found that within the substantial proportion (∼50%) of the transcriptome that has sufficiently long 5’-UTRs to directly observe queueing small ribosomal subunits, 18% show evidence of such queues, which could be indicative of elongation control. Detailed analyses on a smaller number of genes confirm that at least a sub-section of these queues are indicators of translation elongation-limited transcripts. In the absence of any further information, the number of 18% thus puts a tentative upper limit on the proportion of elongation-controlled transcripts in fast growing yeast in general.

Importantly, the statement that 18% of transcripts may be elongation-controlled only applies to the particular growth conditions investigated here. Under stress conditions, the well-studied translational control mechanisms such as Gcn2-mediated phosphorylation of eIF2 (51) or eEF2K-mediated phosphorylation of eEF2 (4) produce a near-complete cessation of translation. If either initiation or elongation activity are strongly reduced, this activity would become rate-limiting for the entire transcriptome, and the distinction between initiation- or elongation-controlled transcripts would then become meaningless. However, as our data show, for moderate regulation the division of the transcriptome into the two pools means that these pools may be separately regulatable, and this appears to occur eg during adapted growth under osmotic stress. Moreover, even during strong translational regulation, the dynamics with which the two pools respond can be different (figure 6).

Following the identification of a potential pool of elongation-controlled mRNAs in yeast we asked whether this regulation was associated with particular pathways. However, GO analyses of the 638 genes showing evidence of a queueing SSU did not reveal any enrichment for particular processes, functions or components (data not shown). This is consistent with the anecdotal evidence from the literature which does not highlight particular functions of elongation controlled mRNAs either, having for example identified components of the molecular clock in *Neurospora crassa* (11), ribosomal proteins in baker’s yeast (12), and proteins with neuronal functions in mammals (6). Thus, it appears that evolution placed particular genes throughout the wider landscape of molecular processes under the control of translation elongation. Further work will be required to elucidate the detailed role of these genes in regulating individual pathways.

## Supporting information

Supplemental file 1: the relatinship between uORFS and queueing subunit footprints.

Experimental details of the ORF replacement procedure.

## Acknowledgements

We are grateful to the Leverhulme Trust (UK) for funding this work through a Research Project Grant (RPG_2014_32, to TVDH). We are grateful for their kind gifts of antibodies to Prof. Yury Chernoff at the Georgia Institute of Technology (Atlanta, USA), Dr. Jordi Tamarit at the Universitat de Lleida (Spain), Prof. Roland Lill at the Universität Marburg (Germany), and Dr. Campbell Gourlay at the University of Kent (UK); and for the gift of a *tef1* deletion strain to Paula Ludovico at the Universidade do Minho (Portugal).

