## Supplemental file 1: the relatinship between uORFS and queueing subunit footprints. for "On translational control by ribosome speed in *S. cerevisiae*"

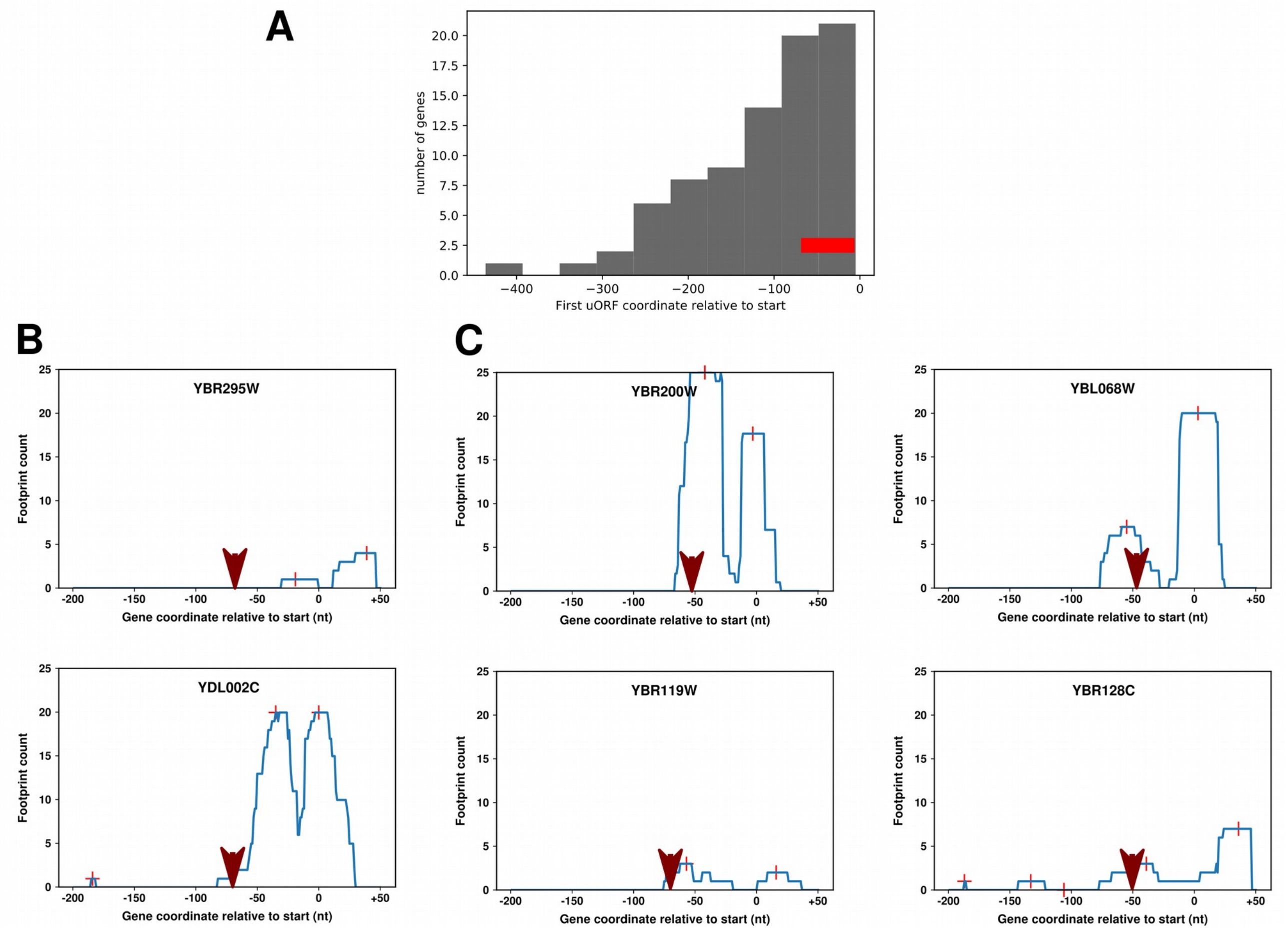


**Supplemental Figure 1. Interaction between apparent queuing small subunit peaks and the presence of upstream open reading frames (uORFs). A,** locations of the most upstream uORF starts in yeast genes annotated as presenting a queuing SSU in our dataset. 35% of uORF containing genes in this set have the uORF start codon in a location overlapping with the -15 to -60 analysis window (indicated by the red bar), ie for these genes the presence of the uORF could confound the queuing SSU analysis. **B, C,** selected SSU footprints for indivdiual uORF-containing genes. Footprinting data are from the dataset by Archer *et al*., and footprint peak locations (crosshairs) were identified in our analyses as described in the main text. Red arrows indicate the location of the 5'-most uORF start codon (ie the uORF start predicted to generate the highest SSU peak through uORF initiation events). **B,** Two examples of genes where the uORF location does not coincide with the second SSU peak, indicating that the second peak is uORF-independent. **C,** four examples of genes where the second SSU peak coincides with thre location of the uORF start, where this second peak may be caused by uORF initiation events rather than ribosome queuing. For YBR119W and YBR200W, the footprint peak on the main AUG is lower than the peak over theuORF start, as would be expected for most uORF-containing genes.
