## Supplementary material for "On translational control by ribosome speed in *S. cerevisiae*": Experimental details of the ORF replacement procedure.

### Construction of gene replacement strains

**1 Sequence design.** An optimised coding sequence was designed by selecting the fastest decoded codon for each amino acid of the protein sequence, according our previous analyses (1). Upstream and downstream sequences surrounding the wild-type gene (60-80 nucleotides) were appended to the 5'- and 3'-ends of the optimised sequence to facilitate homologous recombination. Sequences were synthesised by Genscript (Piscataway, New Jersy, USA) or Eurofins Genomics (Wolverhampton, UK).

**2 guide RNA vector design.** Guide RNA sequences targetting each gene were selected based on ChopChop (2). In each case, we selected the one of the top five scoring guide RNAs located nearest the centre of the target ORF. Paired DNA oligo sequences encoding the guide RNAs were designed and cloned into the dual Cas9/guide RNA expresison vector pML104 as described (3).

**3 transformation.** Optimised genes including flanking chromosomal regions were excised from the synthesis vectors using suitable flanking restriction enzymes or, where no such enzymes were available, were amplified by PCR. 2 µl of pML104 vector containing the gRNA sequences and 0.1-1 µg of linear, optimised gene DNA were co-transformed into yeast strain BY4741 using a standard lithium acetate-based transformation procedure (4) and selection on plates lacking uracil.

**4 confirmation of integration.** For confirmation, primer triplets were designed so that one forward primer annealed around 200 bp upstream of the ORF start, one reverse primer annealed 200 to 300 bp downstream of the ORF start at a sequence specific to the wild-type (non-optimised) gene, and a second primer annealed 300 to 400 bp downstream of the ORF start at a sequence specific to the codon optimised gene. Following colony PCRs containing small amounts of freshly grown yeast cells and all three primers, colonies containing wild-type or optimised genes could be distinguished by the size of the amplification product (increased for the optimised gene). 20 to 200 colonies were screened for each replacement, with targeting efficiencies ranging from 1 in 10 to 1 in 200.

**5 assessing expression levels.** Protein expression levels were determined using specific antibodies sourced mostly from publishing labs (anti-Ade2, anti-Grx5, anti-Nbp35, anti-Ras2, anti-Sup35). Anti-Cdc10 antibody was purchased from Abmart (Shanghai, China, X2-P25342). mRNA expression levels were determined by quantitative real-time PCR, using primers annealing at sites that were identical between the optimised and wild-type genes.

Specific sequences of all DNA-based reagents are given in table 1.

**Table 1. Gene and primer sequences**

| Gene | Optimised Sequence (ENA Acc No) | gRNA oligos | confirmation oligos | qPCR oligos |
|---|---|---|---|---|
| *ADE2* | LT908469 | GATCTGGAAAAGGAGCCATTAACGGTTTTAGA<br>GCTAG,<br>CTAGCTCTAAAACCGTTAATGGCTCCTTTTCCA | GCCGAGAATTTTGTAACACCAACATAACAC,<br>CTCAATCGTTAGCACATCACATTTTTCAGC,<br>GAAGTTTCGGACGCTTGTTCGACC | GAAACTGTCGGTTACGAAGC,<br>TAGGTATATCATTTTATAATTATTTGCTGTA<br>CAAG |
| *CDC10* | LT908470 | GATCTCAAGGCTTCAACGTCAAGGGTTTTAGA<br>GCTAG,<br>CTAGCTCTAAAACCCTTGACGTTGAAGCCTTG<br>A | CAGTAATACTTAACTTTTTTCAGGC,<br>GCGAACGCGGTCCTCCACAAG,<br>CTTTGCGCAGTCAATTCTTTTCTC | TGATTTTGTTACCACTGGAAGCAAAATAG<br>G,<br>TGAGAATACATTACAAATCTTTAAAAAAAT<br>AGCAAATGTAC |
| *GRX5* | LT908471 | GATCGAAGACCCAGAGCTACGTGAGTTTTAGA<br>GCTAG,<br>CTAGCTCTAAAACTCACGTAGCTCTGGGTCTTC | GCGTTCTAGTCCCAAAGGGACAAGAAG,<br>GAAAACTCTTTGATACCTTCACGTAGC,<br>CAATTCACCGGATCTCGCCATGGA | TGATTTTGTTACCACTGGAAGCAAAATAG<br>G,<br>GCAAATGTACATGCATATATAAATATGGAT<br>CGTAAAG |
| *HIS3*[a] | LT908472 | -- | -- | ATGTAGTGACACCGATTATTTA,<br>TACATACTTACTGACATTCATAG |
| *NBP35* | LT908473 | GATCGATTCCGCCATTATATGGAGGTTTTAGAG<br>CTAG,<br>CTAGCTCTAAAACCTCCATATAATGGCGGAATC | GACATAGCACCCTACATAAATAAGTCATCC,<br>CTGTGATCAATGGAATATCTGGATCAGGG,<br>CGCACCGACTTGCAAGTCTTCGTCCG | TAGTATATTTTATCAAAAAGAATAAAAGGA<br>AGTGCATTAGAGG,<br>CGCTTTTGGAGTAATTAGTAGCATTATATAT<br>GG |
| *RAS2* | LT908474 | GATCTGCTAAGCAAGCAATCAACGGTTTTAGA<br>GCTAG,<br>CTAGCTCTAAAACCGTTGATTGCTTGCTTAGCA | GCCGAGAATTTTGTAACACCAACATAACAC,<br>CTCAATCGTTAGCACATCACATTTTTCAGC,<br>GAAGTTTCGGACGCTTGTTCGACC | AGACGAAGGCGGCAAGTACA,<br>TCCTGTGGCCACCGCTATTT |
| *SUP35*[b] | LT908475 | -- | TTATATCTTACATCATCGTATAATATGATCTT,<br>GGCGTCGGGATTGTACTGTTGATAG,<br>CCGCGTCCGGGTTGTATTGTTGGTAA | CTGCCCACTAGCAACAATGTC,<br>CCTTGGTATCTGTTGTTACCTTG |

[a] Construction of the *HIS3* gene replacement strain was performed using a CRISPR independent strategy and is described in Kazana *et al.* 2014

[b] The wild-type and optimised *SUP35* alleles were compared using plasmid expression systems in a shuffling strain containing a chromosomal *SUP35* deletion
